# miRNAs Copy Number Variations repertoire as hallmark indicator of cancer species predisposition

**DOI:** 10.1101/2021.09.21.461294

**Authors:** Chiara Vischioni, Fabio Bove, Federica Mandreoli, Riccardo Martoglia, Valentino Pisi, Cristian Taccioli

**Author notes:** **Materials & Correspondence:** Correspondence and requests for materials should be addressed to C.T.

## Abstract

Aging is one of the hallmarks of multiple human diseases, including cancer. However, the molecular mechanisms associated with high longevity and low cancer incidence percentages characterizing long-living organisms have not been fully understood yet. In this context, we hypothesized that variations in the number of copies (CNVs) of specific genes may protect some species from cancer onset. Based on the statistical comparison of gene copy numbers within the genomes of cancer -prone and -resistant organisms, we identified novel gene targets linked to the tumor predisposition of a species, such as CD52, SAT1 and SUMO protein family members. Furthermore, for the first time, we were able to discover that, considering the entire genome copy number landscape of a species, microRNAs (miRNAs) are among the most significant gene families enriched for cancer progression and predisposition. However, their roles in ageing and cancer resistance from a comparative perspective remains largely unknown. To this end, we identified through bioinformatics analysis, several alterations in miRNAs copy number patterns, represented by duplication of miR-221, miR-222, miR-21, miR-372, miR-30b, miR-30d and miR-31 among others. Therefore, our analysis provides the first evidence that an altered copy number miRNAs signature is able to statistically discriminate species more susceptible to cancer than those that are tumor resistant, helping researchers to discover new possible therapeutic targets involved in tumor predisposition.

---

Aging is one of the hallmarks of cancer insurgence, being considered also one of its possible related risk factors [1]. Therefore, it is probable that, in order to maintain high longevity rate, some species have developed intrinsic molecular mechanisms that protect them from cancer onset or development [2]. Along this assumption, as two binary parallel lines, those organisms that live longer should theoretically possess a higher risk of cancer occurrence. Nevertheless, considering different species, according to Peto’s Paradox theory [3], the body size of an organism and/or its lifespan expectation are not directly correlated with the species percentage of cancer incidence. After more than 40 years of research, the solution to this puzzling paradox is still an open challenge to be solved. Mammals have evolved lifespan and cancer incidence rates which vary among species [4], however mechanisms underlying these differences are still unclear. For example, despite its small size, the naked mole rat, being able to live more than 30 years, is, to date, the longest-living member of the rodent family. Several studies highlighted that, besides the delayed aging, this species also shows the capacity to resist spontaneous and experimentally induced tumorigenesis [5-8]. Conversely, in mice, the cancer-related mortality can reach 90%, coupled with a species maximum life expectancy of four years [9]. With a percentage of cancer incidence of approximately 4.81%, the elephant has been pinpointed as another cancer resistant species [10]. This value is considerably low compared to the human one, for example (approximately 20%) [11]. Interestingly, in 2016, indeed, Sulak and co-authors found that the genome of the African elephant encodes 20 copies of TP53. This amplification of TP53 gene as the “*guardian of the genome stability*”, could be at the basis of its anti-cancer and longevity mechanisms, leading, for example, to increased levels of apoptotic events in response to DNA damage [12]. Indeed, according to Caulin and Maley (2011) [13], the genome of large long-living organisms can reveal an altered number of tumor suppressors and oncogenes (in multiple or reduced copies), that might represent a possible mechanism underlying their capacity of exceeding the threshold of cancer onset, despite their phenotypic predisposition such as size and longevity [13]. Copy Number Variations (CNVs) are duplications or deletions of genomic regions which can be associated with phenotypic alterations, including tumorigenic diseases [14]. In particular, a variation in the gene copy numbers can activate or inactivate tumor suppressors and oncogenes, leading to the development of cancer [15]. Within this framework, long-living animals have to rely on compensatory mechanisms to suppress and prevent cancer progression, that can be straightened by different molecular and genomics mechanisms such as altered gene copy numbers that increase the number of tumor suppressors paralogues or reduce copies of oncogenes [16-17].

In order to test the hypothesis that genomic CNVs are related to the cancer incidence rate of a species, we compared the entire genomic copy number landscape of 9 different mammals (5 cancer resistant and 4 cancer prone organisms), to retrieve among their genomes, which target genes are able to significantly discriminate between these two groups (Table1 & Supplementary Table1-2). A list of the most significant hits (Wilcoxon rank sum test, p-value < 0.05), including known tumor suppressors and oncogenes, is reported in Table1 (See Supplementary Table2 [S2] for the extended version). Our analysis, which exclusively considered the variation in number of gene copies within different species, was able to identify those genes involved in biological processes related to cancer development and maintenance. In this context, the two-group comparison allowed us to identify a clear existing distinction in terms of distribution in the number of gene copies between cancer-prone and cancer-resistant species (Figure1A). Notably, among the most significant genes presenting an altered number of copies we found CD52, SAT1, DMD, EIF5, SUMO2, SUMO3, SUMO4, S100A16, MBD1, MBD2, MBD3, FGFBP1, FOXJ1, NUPR1, SELENOW and JUND. Some of these, such as DMD, MDB1, NUPR1 and JUND have been already well described as tumor suppressors or oncogenes [18-21], whereas the others do not officially belong to any of these two categories, but they have been proposed as key regulators in biological processes such as cell proliferation, migration, and cancer development and progression [22-31]. For instance, CD52 gene (higher number of copies in the cancer prone group), a membrane glycoprotein expressed on the surface of mature lymphocytes, monocytes and dendritic cells [22], resulted as one of the most significant hits of our analysis (p-value = 0.007). Recently, Wang and co-authors [22] identified CD52 as a key role player in tumor immunity, affecting tumor behavior by regulating the associated tumor microenvironment. With the same significant p-value of 0.007, we also identified SAT1 gene (higher number of copies in the cancer prone group) as one of the possible targets to be further investigated in the context of tumor onset. In particular, this gene can regulate and drive brain tumor aggressiveness, promoting molecular pathways in response to DNA damage and regulation of cell cycle [23]. Another significant gene resulting from our analysis was represented by the SUMO protein family members (higher number of copies in the cancer resistant group). During cell cycle progression, many tumor suppressors and oncogenes are regulated via SUMOylation [36], a biological process that, if deregulated, can lead to genome instability and altered cell proliferation [25]. In this context, it is evident that some tumors could be dependent on the functional SUMO pathway, but whether it is required for tumor growth remains to be established. For this reason, SUMO2, SUMO3, and SUMO4 can be potentially exploited in further anti-cancer mechanisms investigations (p-value = 0.024 in the present study), in order to shed light on the regulatory mechanisms underlying the activity of SUMO machinery in an oncogenic framework. Among the most significant hits, we also retrieved some genes that are already known to be tumor suppressor or oncogenes (DMD and JUND respectively). Indeed, mutation or deregulated expression of Duchenne Muscular Dystrophy gene (DMD), is often linked to the development and progression of some major cancer types [18], such as sarcomas, carcinomas, melanomas, lymphomas and brain tumors [37-38], being a well-known tumor suppressor in different types of human cancers. On the other hand, JUND (member of the AP-1 family), that is related to MYC signaling pathway, regulates cell cycle and proliferation, and its overexpression is linked to many types of cancer cell (PCA i.e.) [21]. Thus, as shown in Figure1A, considering the average number of copies of each gene, it is evident that there is a significant difference between the two species categories, which appears even greater if we only refer to the microRNAs CNVs landscape (Figure1B). Therefore, according to our results, important tumor-related miRNAs are able to discriminate between the two organism groups. In particular, miR-424, miR-372, miR-107, miR-124, and miR-1 are just few examples of significant microRNA hits, which possess a suppressor and/or oncogenic role (Figure 1C). For example, miR-424 is known to be a human tumor suppressor that can inhibit cell growth enhancing apoptosis or suppressing cell migration [32]. MiR-372, instead, can participate in WNT cancer molecular pathway [33], whereas the overexpression of miR-107, mediating p53 regulation of hypoxic signaling, can suppress tumor angiogenesis and growth in mice [34]. MiR-1 is another example of tumor suppressor microRNA that has been previously found significantly down-regulated in squamous carcinoma cells [35]. To confirm the hypothesis that microRNAs CNVs can represent one of the most important gene family that can potentially discriminate for cancer predisposition, we also performed an Over-Representation Analysis (ORA) [39] on the total list of significant genes, in order to underlie functional enriched candidates potentially related to cancer (Table2). The most enriched pathways outputted by ORA analysis were: “*MicroRNAs in cancer*”, “*miRNAs involved in DNA damage response*”, “*Metastatic brain tumor*”, “*miRNA targets in ECM and membrane receptors*”, “*let-7 inhibition of ES cell reprogramming*”, and “*miRNAs involvement in the immune response in sepsis*” [40-41]. These results indicate that the most deregulate genes were miRNAs involved in cancer initiation, chronic inflammation, and immune response. Remarkably, performing the ORA analysis applying PANTHER algorithm [42], we also found a significant enrichment in the “*Cadherin signaling network*”, which is a well-known molecular pathway described as key players in cancer [43].

**Table1.** Genomic CNVs landscape comparison. Subset of 25 significant hits resulting from the unpaired 2-group Wilcoxon test (p-value < 0.05). The statistical comparison was made in order to identify those genes able to discriminate between cancer -prone and -resistant species groups, relying exclusively on the genomic copy number values. Some of these genes are already known to be tumor suppressor and/or oncogenes, whereas the others can play pivotal roles in tumorigenesis events, and, for this reason, can be considered as targets to be further investigated and validated in the context of cancer development.

| Gene | p-value <sup>a</sup> | Known_TS <sup>b</sup> | Known_OG <sup>c</sup> | References |
| --- | --- | --- | --- | --- |
| CD52 | 0.007 | NO | NO | [22] |
| SAT1 | 0.007 | NO | NO | [23] |
| MIR424 | 0.009 | YES | NO | [32] |
| MIR372 | 0.010 | NO | YES | [33,51] |
| DMD | 0.014 | YES | NO | [18] |
| EIF5 | 0.017 | NO | NO | [24] |
| MIR107 | 0.022 | YES | YES | [34,66] |
| MIR124-1, MIR124-2, MIR124-3 | 0.022 | YES | NO | [67] |
| SUMO2, SUMO3, SUMO4 | 0.024 | NO | NO | [25-26] |
| MIR506 | 0.029 | YES | YES | [68] |
| MIR509-1 | 0.029 | NO | NO | [69] |
| MIR511 | 0.029 | YES | NO | [70] |
| MIR514A1, MIR514A3, MIR514B | 0.029 | NO | NO | [71] |
| MIR378A | 0.030 | YES | NO | [72] |
| S100A16 | 0.030 | NO | NO | [27] |
| MBD1, MBD2, MBD3 | 0.031 | NO | YES (MDB1) | [19] |
| FGFBP1 | 0.032 | NO | NO | [28] |
| FOXJ1 | 0.032 | NO | NO | [29] |
| MIR1-1, MIR1-2 | 0.032 | YES | NO | [35,73] |
| MIR206 | 0.032 | YES | NO | [74] |
| MIR340 | 0.032 | YES | NO | [75] |
| MIR542 | 0.032 | NO | NO | [76] |
| NUPR1 | 0.032 | YES | NO | [20] |
| SELENOW | 0.032 | NO | NO | [30-31] |
| JUND | 0.034 | NO | YES | [21] |

<sup>a</sup> Unpaired 2-group Wilcoxon test
<sup>b</sup> https://bioinfo.uth.edu/TSGene (Zhao, M., Kim, P., Mitra, R., Zhao, J., & Zhao, Z. (2016). TSGene 2.0: an updated literature-based knowledgebase for tumor suppressor genes. Nucleic acids research, 44(D1), D1023-D1031)
<sup>c</sup> http://ongene.bioinfo-minzhao.org (Liu, Y., Sun, J., & Zhao, M. (2017). ONGene: a literature-based database for human oncogenes)

**Figure1.**
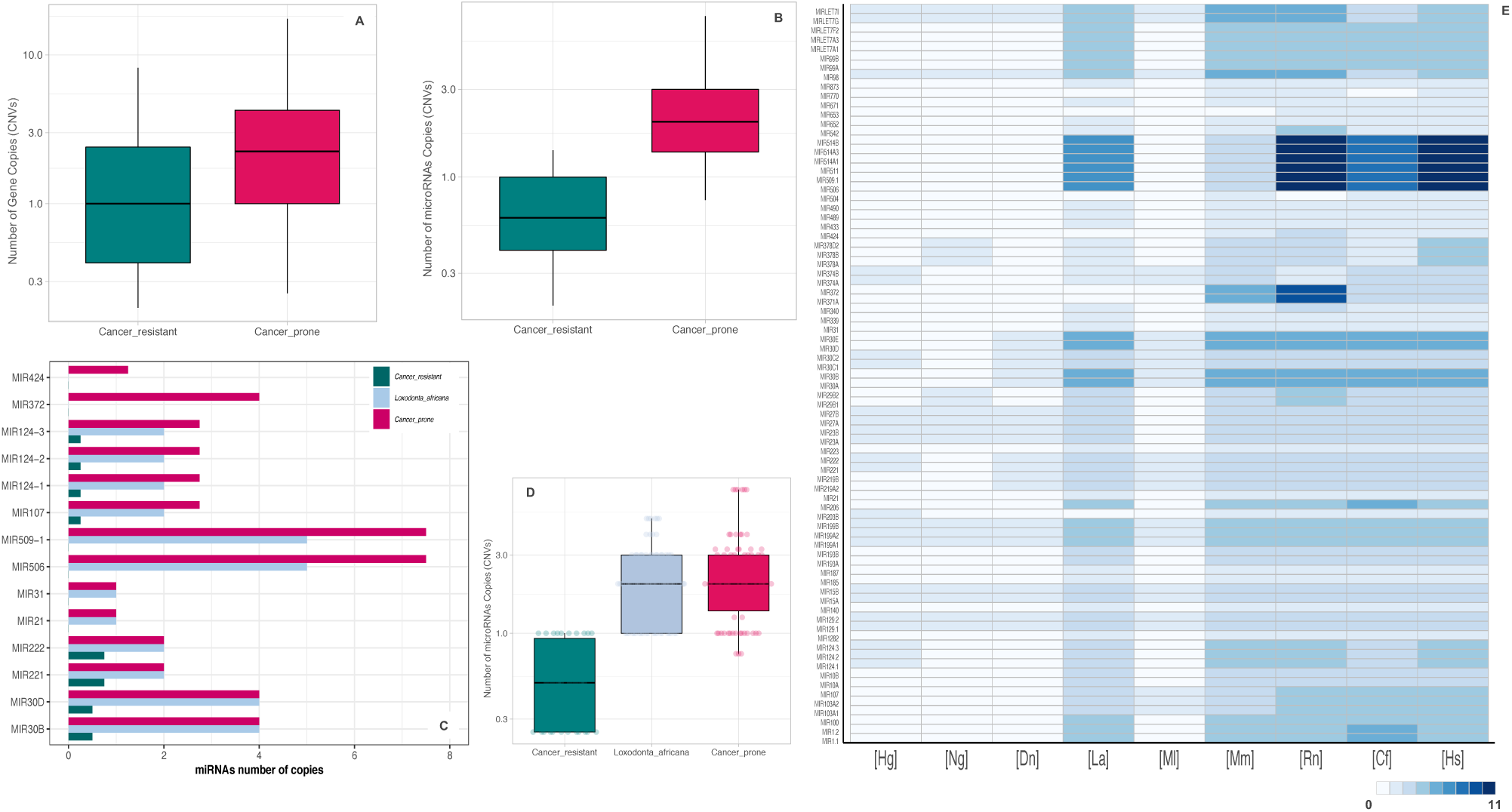
CNVs landscapes comparisons: A, Boxplot of the distribution of significant genes CNVs in cancer prone *vs* cancer resistant species. **B**, Boxplot of the distribution of significant microRNAs CNVs in cancer prone *vs* cancer resistant species. **C**,**D** Bar- and Box-plot of significant microRNAs CNVs in cancer prone species, cancer resistant species, and *Loxodonta Africana*. **C** highlights that the microRNAs repertoire of *Loxodonta africana* seems to reflect the cancer prone miRNAs copy number alteration landscape, rather than the one typical of the cancer resistant organisms. In **D**, each dot corresponds to the average of the number of copy of every significant miRNA. Cancer resistant species are highlighted in green, cancer prone species in red, whereas *Loxodonta Africana* in light violet. In the boxplots, the *Y*-axis scale has been changed to log one. The boxplots are built considering the average number of copies of each gene in the two different target groups. **E**, heatmap representing the MicroRNAs CNVs repertoire within the 9 analyzed species. The color code corresponds to the gene number of copies. [Hg]: *Heterocephalus glaber*; [Ng]: *Nannospalax galili*; [Dn]: *Dasypus novemcinctus*; [La]: *Loxodonta Africana*; [Ml]: *Myotis lucifugus*; [Mm]: *Mus musculus*; [Rr]: *Rattus norvegicus*; [Cf]: *Canis familiaris*; [Hs]: *Homo sapiens*. [Hg], [Ng], [Dn], [La] and [Mm] are considered “cancer resistant” organisms, having a cancer incidence rate lower than 5%. [Mm], [Rr], [Cf] and [Hs] are considered “cancer prone” species, given their percentage of cancer rate higher than 5%.

**Table2.** Pathway analysis. Gene Over-Representation Analysis (ORA) using KEGG, PANTHER, and Wikipathway databases. This statistical test for overrepresentation determines whether a given gene is found statistically more (or less) often in the input list than expected by chance, considering the representation of the pathway genes in the entire reference genome. ORA was performed by the WebGesTalt functional enrichment analysis tool available at http://www.webgestalt.org. The enrichment test used Benjamini-Hochberg’s FDR correction (FDR < 0.05). CNVs data were previously analyzed by an unpaired 2-group Wilcoxon test (p-value < 0.05). Significant genes altered in their number of copies within the entire genomic landscape were used to perform the ORA analysis, which highlighted a significant enrichment in MicroRNAs and cancer related pathways.

|  | Description | FDR (BH) | Genes |
| --- | --- | --- | --- |
| **KEGG** | MicroRNAs in cancer | 0 | MIR103A1; MIR103A2; MIR107; MIR124-1; MIR124-2; MIR124-3; MIR1-1; MIR1-2; MIR206; MIR100; MIR10A; MIR10B; MIR129-1; MIR129-2; MIR15A; MIR15B; MIR193B; MIR199A1; MIR199A2; MIR199B; MIR203B; MIR21; MIR223; MIR31; MIR99A; MIRLET7A1; MIRLET7A3; MIRLET7F2; MIR29B1; MIR29B2; MIRLET7G; MIRLET7I; MIR221; MIR222; MIR23A; MIR23B; MIR27A; MIR27B; MIR30C1; MIR30C2; MIR30A; MIR30B; MIR30D; MIR30E. |
|  | Taste transduction | 3.16E-10 | TAS2R10; TAS2R13; TAS2R14; TAS2R19; TAS2R20; TAS2R3; TAS2R30; TAS2R31; TAS2R42; TAS2R43; TAS2R45; TAS2R46; TAS2R50; TAS2R7; TAS2R8; TAS2R9 |
|  | Progesterone-mediated oocyte maturation | 2.43E-04 | SPDYE1; SPDYE11; SPDYE16; SPDYE17; SPDYE2; SPDYE2B; SPDYE3; SPDYE4; SPDYE5; SPDYE6; INS |
|  | Oocyte meiosis | 2.73E-04 | PPP3R2; SPDYE1; SPDYE11; SPDYE16; SPDYE17; SPDYE2; SPDYE2B; SPDYE3; SPDYE4; SPDYE5; SPDYE6; INS |
| **PANTHER** | Cadherin signaling pathway | 0.040196 | PCDHB14; PCDHB7; PCDHGB1; PCDHB16; PCDHB6; PCDHGB4; PCDHGA6; PCDHGB6; PCDHGB7 |
| **Wikipathway** | miRNAs involved in DNA damage response | 3.76E-09 | MIR371A; MIR372; MIR542; MIR100; MIR15B; MIRLET7A1; MIR374B; MIR221; MIR222; MIR23A; MIR23B; MIR27A; MIR27B |
|  | Alzheimers Disease | 5.31E-05 | MIR124-1; MIR124-2; MIR124-3; MIR10A; MIR129-1; MIR129-2; MIR199B; MIR21; MIR433; MIR671; MIR873; PPP3R2; MIR29B1; MIR30C2; MIR219A2 |
|  | Metastatic brain tumor | 0.00230738 | MIRLET7A1; MIRLET7A3; MIRLET7F2; MIR29B1; MIR29B2; MIRLET7G |
|  | miRNA targets in ECM and membrane receptors | 0.00230738 | MIR107; MIR15B; MIR30C1; MIR30C2; MIR30B; MIR30D; MIR30E |
|  | MicroRNAs in cardiomyocyte hypertrophy | 0.00276823 | MIR103A1; MIR103A2; MIR140; MIR15B; MIR185; MIR199A1; MIR199A2; MIR23A; MIR27B; MIR30E |
|  | Cell Differentiation - Index | 0.012506063 | MIR1-1; MIR206; MIR199A1; MIR199A2; MIR221; MIR222 |
|  | let-7 inhibition of ES cell reprogramming | 0.01250606 | MIRLET7A1; MIRLET7F2; MIRLET7G; MIRLET7I |
|  | miRNAs involvement in the immune response in sepsis | 0.01427985 | MIR187; MIR199A1; MIR199A2; MIR203B; MIR223; MIR29B1; MIRLET7I |
|  | Cell Differentiation - Index expanded | 0.02382508 | MIR1-1; MIR206; MIR199A1; MIR199A2; MIR221; MIR222 |
|  | Role of Osx and miRNAs in tooth development | 0.03346077 | MIRLET7A1; MIRLET7F2; MIR29B1; MIRLET7G; MIRLET7I |

## Cancer related MicroRNAs pathways are among the most significantly enriched biological families

Our results show that miRNAs are the most enriched gene family in discriminating between cancer-prone and cancer-resistant species. The specific role of these miRNAs is not yet fully understood, but we speculate that some of them possess important regulatory functions aimed at defending some species (big size and long lifespan organisms) from cancer, while, at the same time, they are capable of exposing others to tumorigenesis (small size and short lifespan mammals). MicroRNAs (miRNAs) are small post-transcriptional molecular regulators, which are able to modify gene expression levels increasing the amount of mRNA degradation or inhibiting protein translation [44]. Since each single miRNA can regulate the expression of dozens of genes, many authors were able to correlate their activity with cell development, homeostasis and disease [45], including cancer [46-47]. Indeed, some tumorigenic events are caused by a malfunction in the heterogeneous regulatory activity of microRNAs inside the eukaryotic cells. Depending on the specific tissue and on the relationship with the immune system, they can behave both as tumor suppressors and as oncogenes [48]. Furthermore, epigenetic factors and species genetic predisposition can drive their double side behavior, although some of them are evolutionary conserved within vertebrate taxonomic families [49]. Several miRNAs have been already previously described in literature as oncogenes and tumor suppressors. For example, MiR-30b and miR-30d are considered suppressors in those tumors that do not involve immune cells, whereas they have been found upregulated in melanoma [50]. Similarly, for the first time, our analysis revealed several miRNAs candidates that might be involved in a mammalian species cancer predisposition (Figure 1E). Notably, all the miRNAs we have found show many more copies in the cancer prone group compared to the cancer free species (Supplementary table 2 [S2]), and most of them are well known as oncogenes (e.g. miR-221, miR222, and miR-372, etc.). MiR-372, for instance, is not present in cancer free species, whereas it shows multiple copies in almost all those ones belonging to the cancer prone group. This microRNA can play a pivotal role in the initiation of breast cancer and may activate WNT and E2F1 pathway during the epithelial-mesenchymal transition process [33,51]. We also found an amplification in the cancer prone category for miR-221 and miR-222. Extensive literature has described these two RNAs as oncogenes, being deregulated in primary brain tumors and in Acute Lymphoid Leukemia among other malignancies [52-53]. According to our results, surprisingly, cancer prone species showed the amplification of miR-15 and miR-16 tumor suppressors, which are known to be able to regulate cancer proliferation initiation by targeting BCL2 gene [54]. Our hypothesis is that this apparent paradox may underlie a defensive role of these two microRNAs in those species that are, a priori, susceptible to tumor insurgence. Indeed, according to the so-called “*gene dosage hypothesis*”, gains or losses of specific gene copies can have a dramatic impact on the fitness of a species, leading to altered phenotypes due to the change in the expression levels of the affected genes [55]. On the other hand, oncogenes amplification or tumor suppressors deletions are not always detrimental, but can recapitulate tumorigenic events, being drivers or modulators of the disease [56].

Interestingly, compared to the other mammals, elephant miRNAs amplification signature resembles the organisms of cancer prone group (Figure1C-D). In fact, it showed an alteration in the copy numbers of known oncogenes, such as miR-221 and miR-222, together with miR-30b/d and miR-31. This can be due to the species cancer incidence rate, which is proximal to our value of cancer prone threshold (∼5%). In our opinion, *Loxodonta africana*, should be placed in a new category of organisms, which share both oncogenic and cancer free characteristics. In this context, during the evolution, elephants may have selected molecular defenses, such as the amplification of TP53 and pseudogenes [12], with the aim to defend its cells from the tumorigenic action of a high percentage of onco-miRNAs copy number amplification and high longevity. In support of the hypothesis described above, recently, Vazquez and Lynch (2021) [57] reported that, within the Afrotheria order, the tumor suppressor genes found in an altered number of copies was relatively lower compared to what might be expected. This finding can mirror the trade-off mechanism that natural selection has developed during evolution in order to compensate for the multi copies effect which can lead to an increased risk of cancer, due to the unbalanced number of copies of the same genes. On the other hand, cancer-prone organisms (mouse, rat, etc.) included in our analysis, do not develop these gene defenses because they have a lower lifespan, which does not make them particularly exposed to a severe lack of fitness due to cancer progression (except the case of *Homo sapiens* that has reached a high lifespan only recently, thanks to the advance of medicine treatments and health care).

## Conclusions

Being theoretically more susceptible to cancer, big and long living species need additional cancer defense molecular mechanisms. On the other hand, short living and small size organisms might not need them because of their lower intrinsic predisposition to cancer due to their short lifespan rate. CNVs can therefore be considered only one of the multiple protection ways against tumor insurgence that can explain Peto’s paradox. In fact, we hypothesize that, all cancer resistant organisms implemented a series of molecular mechanisms aimed to preserve themselves from their cancer predisposition, which in turn depends on and derives from their own specific evolutionary history. We believe that CNVs that increase the onco-suppressive capacity of specific genes such as microRNAs, can be one of the excellent defenses against tumor diseases in species at risk. Our analysis shows an enrichment of onco-miRNAs, as particularly amplified in cancer prone species group. The specific role of these miRNAs is not totally understood yet, but we think that, some of them, possess important regulatory molecular functions within each species, but simultaneously expose some organisms to the danger of cancer initiation. In our opinion, studying microRNAs that are related to human malignancies from a comparative perspective, can provide additional clues about their role, and potentially point towards novel targets involved in tumorigenic diseases. Focusing on patterns of miRNA copy number changes may, for the first time, lend new insight into the conserved molecular pathways across those species, and leading to the discovery of novel therapeutic approaches.

## Material and Methods

According to the hypothesis that positively selected CNVs tend to recur during cancer progression [58-59], but also during the evolution [60], we have recently developed VarNuCopy database (freely available at http://isgroup.mat.unimore.it:8083/), a tool unique of its kind, which collects the CNVs landscape for multiple organisms, with the aim to compare patterns of copy number changes across the genome of different species. We used a homemade script written in Perl 5.14 and Python 3 in order to download the CNV data from Ensembl comparative genomics resources (http://www.ensembl.org) [61], an ideal system to perform and support vertebrate comparative genomic analyses, given the consistency of gene annotation across the genomes of different vertebrate species. In this context, we leveraged Ensembl’s “gene gain/loss tree” feature, which maps the number of copies of extant homologous gene for each species, as a taxonomic tree view [62]. The Perl API script provided by the Ensembl website was used to access the genomic databases and was used to download all the available CNVs data. We encoded a new homemade Python script in order to format the CNVs data counts as a readable tab delimited matrix, useful to perform the subsequent analysis. Then, using a comparative approach, we analyzed the variation landscape of genes copies among the genome of 9 organisms sub-set in two categories: “cancer resistant” (*Heterocephalus glaber* [Hg], *Nannospalax galili* [Ng], *Dasypus novemcinctus* [Dn], *Loxodonta Africana* [La], *Myotis lucifugus* [Ml]), and “cancer prone” (*Mus musculus* [Mm], *Rattus norvegicus* [Rn], *Canis familiaris* [Cf], and *Homo sapiens* [Hs]) species. We classified as “cancer resistant” those species that have a cancer incidence rate less or equal to 5%. Conversely, “cancer prone” organisms, were referred to those species for which the percentage of tumors found in a certain number of necropsies is higher than 5% (Supplementary Table1 [S1]). We performed a statistical comparison between the CNVs of the two different species groups, cancer prone- and resistant organisms, with the aim to identify new possible gene targets able to discriminate between the two categories. Thus, a statistical unpaired 2-group Wilcoxon test was performed using R.3.1.1 (https://www.r-project.org/), to compare their entire CNVs spectra. To determine if CNVs are enriched in specific gene families, we used Gene SeT AnaLysis Toolkit, a tool for the interpretation of lists of interesting genes which is commonly used to extract biological insights from targets of interest [39]. The set of significant genes were tested for pathway associations using the hyper-geometric test for over-representation analysis (ORA) [63] (Supplementary Table3 [S3]). We selected different pathway enrichment categories (KEGG: https://www.genome.jp/; Wikipathway: https://www.wikipathways.org; Reactome: https://reactome.org/; PANTHER: http://www.pantherdb.org/), considering over-represented those molecular networks with FDR significance level lower than 0.05, after a correction with Benjamini–Hochberg method. In this context, the ORA analysis was the preferred option among the others (e.g. gene set enrichment or network topology-based analysis) in order to obtain biological information underlying the significantly enriched genes, resulting in a reduction in the complexity of the data interpretation [63]. Data processing, plots, and statistical tests were performed with R.3.1.1 (www.cran.r-project.org) and RStudio 1.4.1717 (https://www.rstudio.com/). Figures were made using the ggplot2 R package, in association with different R Shiny apps such as BoxPlotR and PlotsOfData [64-65].

## Data Availability Statement

All data necessary for confirming the conclusions of the article are present within the article, figures, tables, and its supplementary material.

## Acknowledgements

This study was supported by the University of Padova through M.A.P.S. department under the program BIRD213010. We thank Dr. Nicoletta Bianchi for critical reading of our manuscript and for her precious suggestions.

## Author contributions

C.V., C.T., conceived, designed, and performed the research. F.B., F.M., R.M., V.P., established the bioinformatical resource used to generate the data. All authors analyzed the data. C.V. interpreted the data. C.T., F.M., R.M., supervised the project. C.V., C.T wrote the manuscript. All authors have read and agreed to this version of the manuscript.

